# RBAtools: a programming interface for Resource Balance Analysis modelling

**DOI:** 10.1101/2022.12.19.521060

**Authors:** Oliver Bodeit, Inès Ben Samir, Jonathan R. Karr, Anne Goelzer, Wolfram Liebermeister

## Abstract

**Motivation:** Efficient resource allocation contributes to an organism’s fitness and improves success in competition. The Resource Balance Analysis (RBA) computational framework enables the analysis of an organism’s growth-optimal configurations in various environments, at genome-scale. The existing RBApy software enables the construction of RBA models on genome-scale and the calculation of medium-specific, growth-optimal cell states, including metabolic fluxes and the abundance of macromolecular machines. However, to address the needs of non-expert users, there is a need for a simple programming API, easy to use and interoperable with other software through convenient formats for models and data.

**Results:** The RBAtools python package enables the convenient use of RBA models and addresses non-expert users.

As a flexible programming interface, it enables the implementation of custom workflows and simplifies the modification of existing genome-scale RBA models and data export to various formats. The features comprise simulation, model fitting, parameter screens, sensitivity analysis, variability analysis, and the construction of Pareto fronts. Models and data are represented as structured tables, in HTML, and common formats for fluxomics and proteomics visualization.

**Availability:** Details about RBA can be found at rba.inrae.fr. RBAtools documentation, installation instructions, and tutorials are available at sysbioinra.github.io/rbatools.

## Introduction

How can we understand and anticipate the impact of genomic modifications and environmental perturbations on the behavior of microbial cells? A guiding idea is that organisms efficiently allocate their resources (1), (2), (3) to succeed within their ecological niche. Resource Balance Analysis (RBA) is a conceptual and computational framework that implements this principle as a constraint-based modeling method (4), predicting growth-optimal cellular states (2), (5). It allows for simulating responses to genomic (e.g. gene-knockouts and the addition of heterologous metabolic pathways) or environmental perturbations (e.g. nutrient limitation). Due to its formulation as a linear optimization problem (LP), RBA can handle detailed cell models at genome-scale. Available RBApy (6) software facilitates the construction of genome-scale RBA models and the prediction of growth rates and corresponding cellular states (i.e. metabolic fluxes and abundance of molecular machines). However, RBApy lacks functionality for custom analysis workflows and for exploring resource allocation beyond growth-optimal states.

### RBAtools

RBAtools is a programming interface based on RBApy with extended capabilities for the exploration, manipulation and simulation of RBA models. It provides new and more convenient functionality for defining the growth medium, manipulating model components and parameters, and directly editing the LP problem. The new methods can be easily combined to define custom workflows for simulation and analysis. RBAtools also facilitates programmatic access to model components and their relationships, and the export into human-readable formats such as SBtab (7) or CSV. Furthermore, predicted fluxes and protein levels can be converted into intuitive visualizations with Escher maps (8) and Proteomaps (9). RBAtools provides elementary functions for setting parameters or manipulating and solving the LP problem, enabling the implementation of custom algorithms. In addition, it includes high-level methods for simulation and analysis workflows used in resource allocation modeling. Below we showcase some analysis methods from the RBAtools library (for implementation details, see the supplementary materials).

#### Prediction of cell phenotypes for biotechnology

RBAtools provides various modeling methods for biotechnology, to improve the production of added-value compounds: models can be modified to simulate gene knock-outs, enzyme inhibition, enzyme overexpression or underexpression, changes in cell dry-mass composition, and the addition of heterologous metabolic pathways. Known characteristics such as measured metabolite exchanges (e.g added-value compound) or machinery abundances can be imposed, and the resulting phenotype (maximum growth rate, metabolic activity and quantitative proteome) can be inferred. It is possible to predict Monod curves (Fig. 1a) and to infer the minimum required concentration of a limiting substrate in the medium for a fixed given growth rate (as in chemostat).

**Fig. 1.**
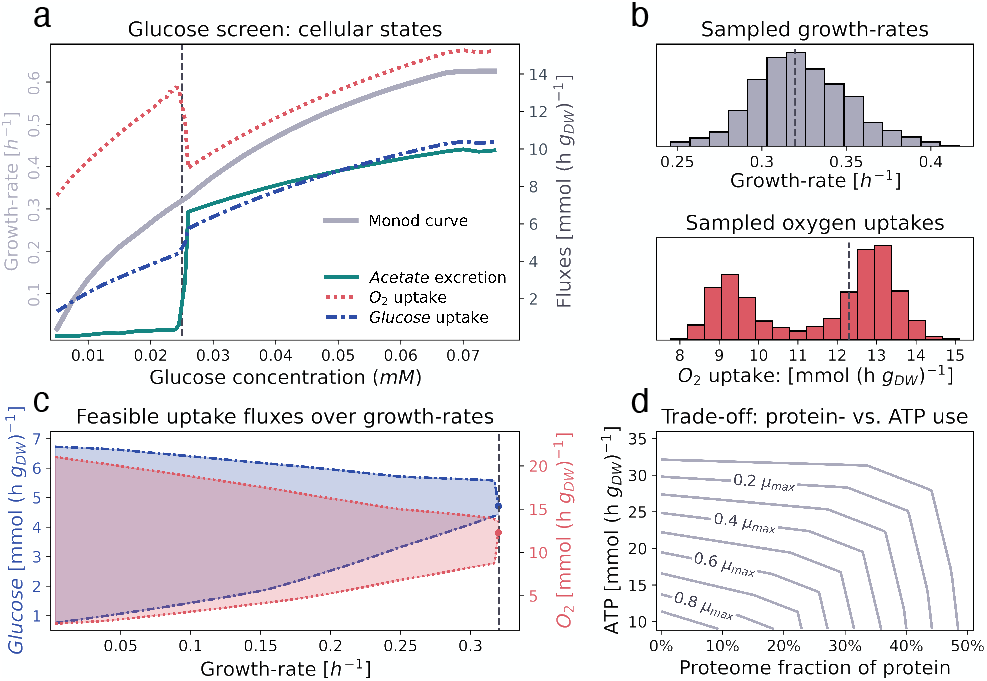
Analyses of a genome-scale *B. subtilis* cell model, performed with RBAtools. (a): Phenotype prediction. Screen over external glucose concentrations as sole carbon source and resulting maximal growth rates (Monod curve) and corresponding optimal exchange fluxes of glucose, oxygen and acetate. A glucose concentration of 0.025 mM, at the onset of overflow metabolism, is used as a reference condition in other panels (dashed black lines). (b): Sensitivity analysis. Global sensitivity of (growth optimal) cellular states to the uncertainty of enzyme efficiencies. For each enzyme efficiency in the model, a scaling factor *x* is applied, where *ln*(*x*) is drawn from a normal distribution with *μ* = 0 and *σ* = ln(1.1). The respective maximal growth rates and optimal oxygen exchanges are determined for 1000 samples. The two modes in the distribution of oxygen uptake rates correspond to two different metabolic configurations, respiration and overflow metabolism, which are normally used at low or high glucose concentrations, respectively. Around the reference concentration of 0.25 mM (critical concentration for wild type), a cell may show either configuration, depending on its enzyme efficiencies, and uncertainty in these parameters may have large effects. (c): Resource Variability Analysis. feasible ranges of glucose and oxygen uptake at growth rates between 0 and the maximum. Around the maximum growth rate the feasible regions collapse to the respective optimal values. (d): Pareto analysis. Pareto efficient trade-offs between investment in non-native cytosolic protein, and additional ATP-turnover at various fractions of the maximal growth rate (*μ*max). Lines represent simulated Pareto fronts.

#### Sensitivity to model parameters

Modifiable model parameters allow for different types of sensitivity analyses, implemented in the RBAtools library. (i) By screening parameter values and predicting associated cellular states, effects on the phenotype can be assessed. (ii) Local fitness sensitivities, defined as the partial derivative of growth rate to the parameter value, can be determined. (iii) Global parameter uncertainty and the resulting variability can be studied by sampling global sets of enzyme capacities and predicting the resulting phenotypes (Fig. 1b).

#### Resource Variability Analysis and multi-objective optimization

Biotechnology needs to consider trade-offs between cell variables (e.g. between cell growth and the production of compound production). Similar to Flux Variability Analysis (10), RBAtools allows users to determine feasible ranges of metabolic fluxes and machinery concentrations, providing ultimate bounds at predefined growth rates (Fig. 1c). By relating these ranges to growth rate, tradeoffs between extreme production/consumption capabilities and cellular fitness can be assessed. RBAtools can also directly trace Pareto fronts between state variables (e.g. variables representing the usage of alternative metabolic routes, or the production of additional ATP versus additional proteins) at a given growth rate and environment (Fig. 1d). Aside from these predefined procedures, versed users can easily implement their own custom methods based on more fundamental functionalities on model- and LP manipulation.

## Conclusion

We developed a versatile and convenient programming interface for RBA models, more flexible and user-friendly than existing tools, enabling deep exploration of cellular behavior on genome-scale. It leverages the idea of cellular resource allocation to model the physiology of cells by combining biochemical facts, optimality considerations, and organism-specific empirical knowledge. While RBApy remains the tool for building RBA models, RBAtools with its simplified interface and extended functionality allows users to easily simulate the impact of genomic or environmental perturbations on cell phenotypes. This makes it useful for a wide range of applications in synthetic biology, metabolic engineering or white biotechnology.

## Acknowledgments and Funding

This work was supported by the German Research Foundation (Ll 1676/2-2) and the National Institutes of Health awards P41EB023912 and R35GM119771.

We thank Ana Bulović, Stephan Fischer, Vincent Fromion, Marc Dinh, Edda Klipp, Hermann-Georg Holzhütter, Oliver Ebenhöh and Christian Poirier for helpful discussions and support.

## Article template

We thank Ricardo Henriques for providing us and the community with the LaTex template for publications on bioRxiv, we used to write this manuscript.

